## Supplementary Material for "Genomic and local microenvironment effects shaping epithelial-to-mesenchymal trajectories in cancer"

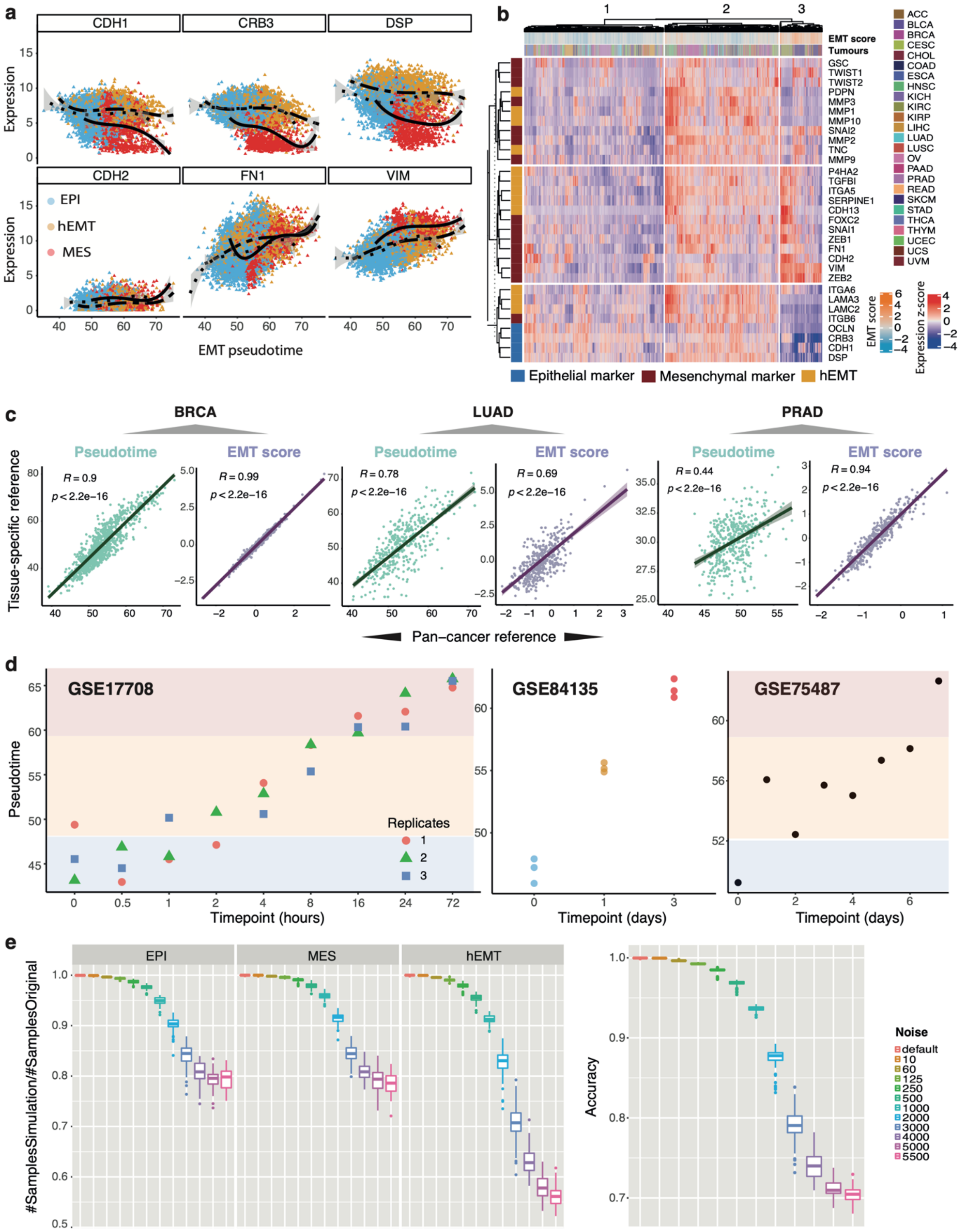


**Supplementary Figure S1. Validation of EMT trajectory reconstruction methodology**. (a) Expression of epithelial/mesenchymal markers along the EMT pseudotime derived from single cell data. Each dot represents a TCGA sample and is coloured according to the assigned EMT state. (b) Heat map highlighting the pan-cancer expression of known EMT markers (coloured by their state-specificity). (c) Correlation between pseudotime estimates (green) and EMT scores (purple) obtained from a pan-cancer versus tissue-specific consensus single cell reference, shown for four independent cancer tissues. The x axis depicts the estimates calculated with a pan-cancer reference, and the y axis with a tissue-specific reference. (d) Application of the EMT trajectory reconstruction method in three longitudinal datasets: GSE17708, a time course experiment of A549 lung adenocarcinoma lines treated with TGF-beta; GSE84135, a time course EMT transition experiment in hSAEC airway epithelial cells; and GSE75487, 7 day EMT transformation of H358 non-small cell lung cancer cells under doxycycline treatment to induce Zeb1. The pseudotime estimate increases with time as expected for gradually transforming cells. (e) Left panel: Fraction of samples correctly assigned to the original HMM state with increasing levels of gene expression noise in the original data. Right panel: Accuracy of predicting the original HMM states with increasing levels of gene expression noise in the original data.

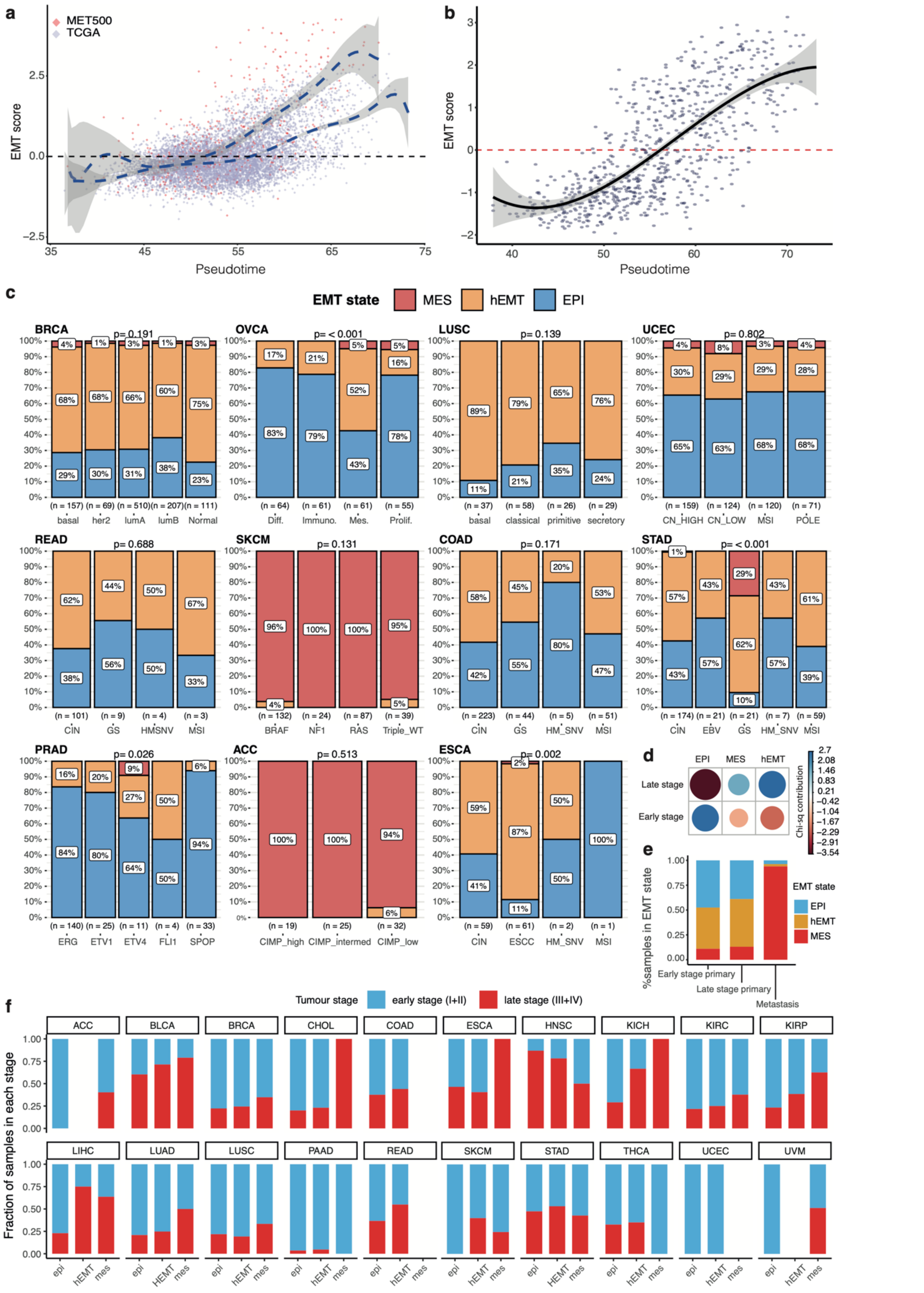

**Supplementary Figure S2.** (a) EMT scores of the MET500 (red points) and TCGA (blue points) samples plotted along the EMT pseudotime derived from single cell data. (b) EMT scores of the CCLE samples plotted along the EMT pseudotime. (c) EMT state distribution by molecular cancer subtypes. (d) Relation between the EMT states and the clinical cancer stage, computed using Chi-square statistics. The circles represent the associations between EMT states and cancer stage (early/late). The colour and size define the strength of the association. (e) EMT score distribution compared between early, late stage primaries and metastatic samples. (f) Tumour stage distribution across cancer types and EMT macro-states.

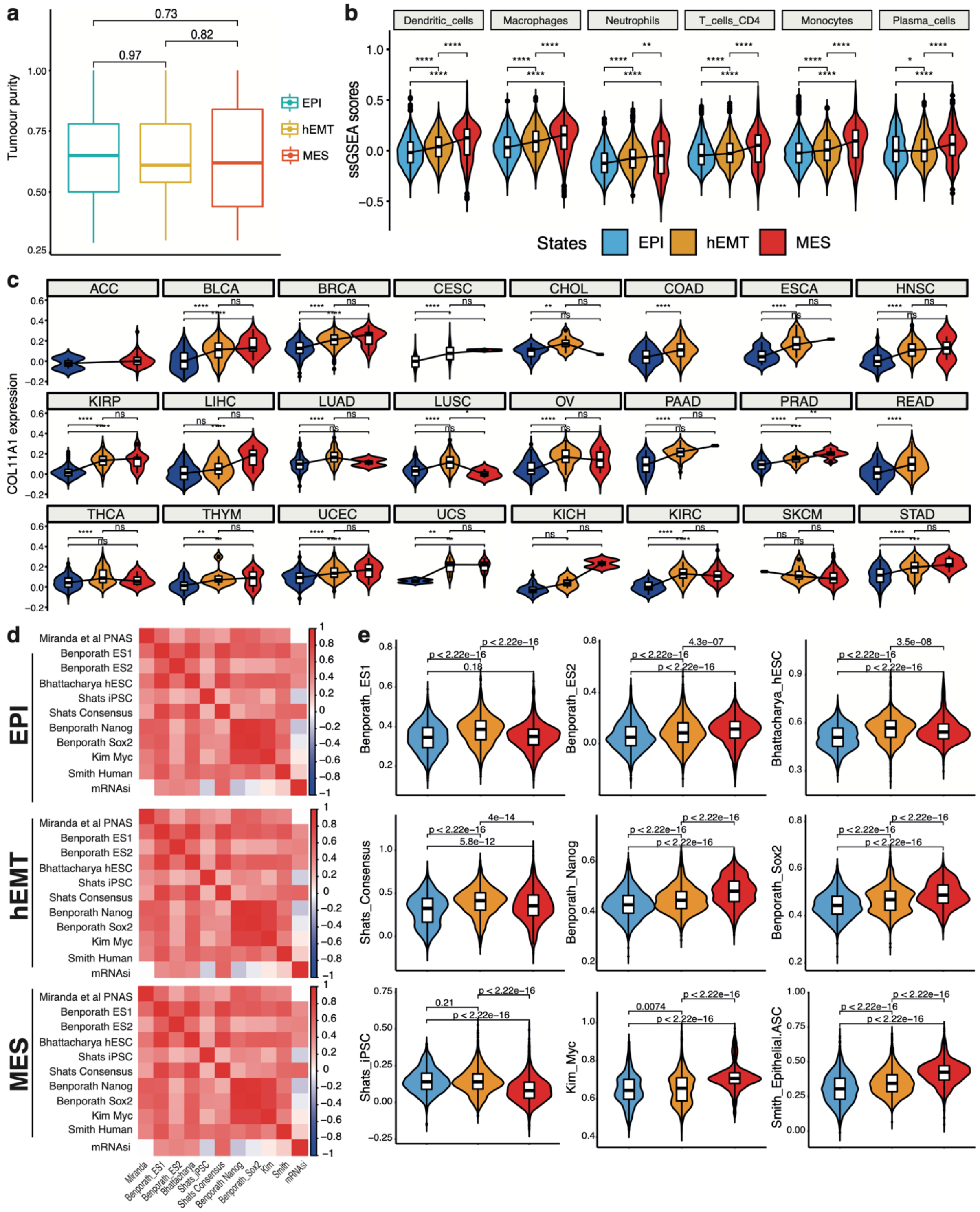


**Supplementary Figure S3. Tumour extrinsic properties in relation to EMT.** (a) Tumour purity compared between the three discrete EMT macro-states pan-cancer. (b) Tumour microenvironment composition compared between the three EMT macro-states. (c) Active fibroblasts infiltration levels (quantified based on COL11A1 expression) compared between the three EMT macro-states, by cancer type. (d) Correlation heat maps of the stemness scores assessed in this study. The strength of the Pearson correlation is highlighted by the colour gradient. (e) Stemness scores compared between each EMT state. *p<0.05; **p<0.01; ***p<0.0001; ****p<0.00001.

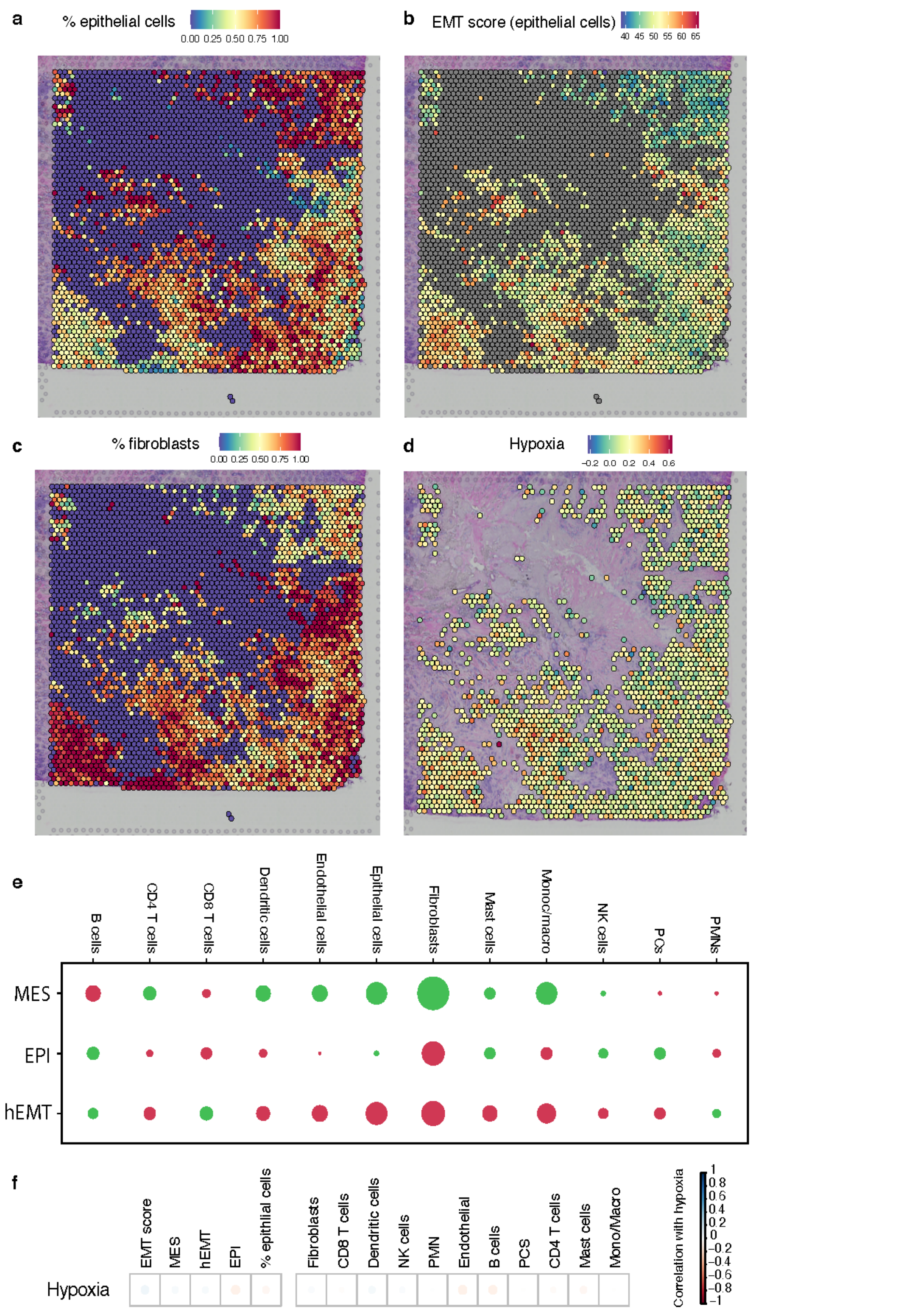


**Supplementary Figure S4. Spatial patterns of EMT in Patient 3.** (a-d) Spot annotations of the fraction of epithelial cells (a), EMT scores across these epithelial spots (b), fraction of fibroblasts (c) and hypoxia (d) within individual spots profiled across the tissue within the breast cancer slide of Patient 3. The blue to red gradient indicates increased expression of markers of the specific cell state or increased fraction of cell types. (e) Enrichment (green) and depletion (red) of cell types in each EMT-based cluster identified within the slide. The plots represent the difference between the average cell type proportion value per region, compared to a permuted spot value (calculated 10,000 times). The plot marker size corresponds to the absolute enrichment score, and the colour represents the enrichment sign (red for negative and green for positive). (f) Correlation between hypoxia and individual cell types and states. Blue indicates positive correlation, red indicates negative correlation, with the circle size being proportional to the correlation value. The correlations with all cell types are relatively weak.

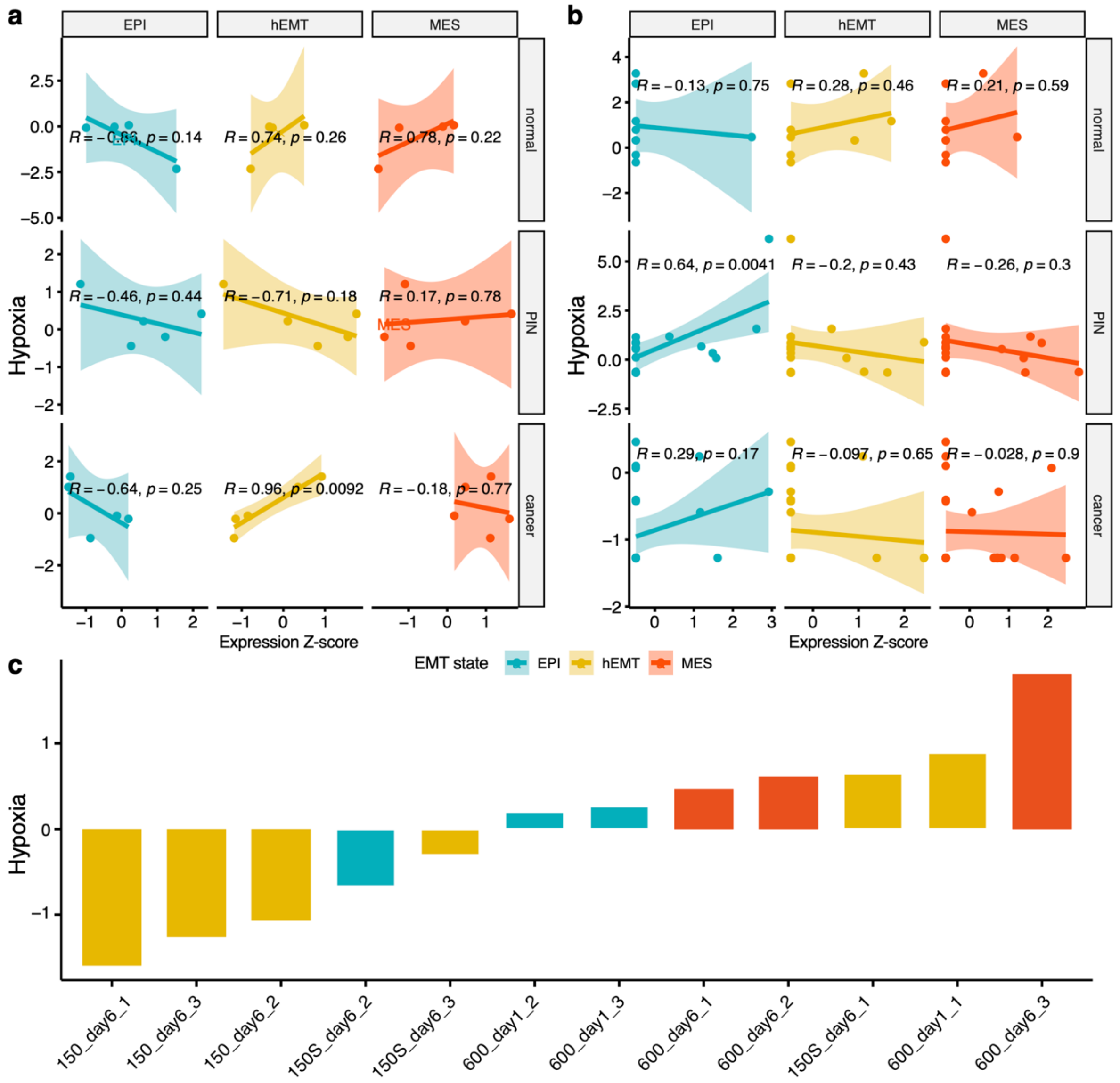


**Supplementary Figure S5. Relation between hypoxia and EMT transformation in external datasets.** (a) Correlations between hypoxia and EPI, hEMT and MES enrichment scores, respectively, measured in spatially profiled normal, PIN and prostate cancer samples from tissue section 1.2 from Berglund et al (Nat Commun 2018). (b) Correlations between hypoxia and EPI, hEMT and MES enrichment scores, respectively, measured in spatially profiled normal, PIN and prostate cancer samples from tissue section 3.3 from Berglund et al (Nat Commun 2018). (c) Hypoxia scores measured in a 3D microtumour model of breast cancer at different days after inducing cellular migration. Bars are coloured according to the assigned EMT macro-state for the respective sample, split by quartiles of expression.

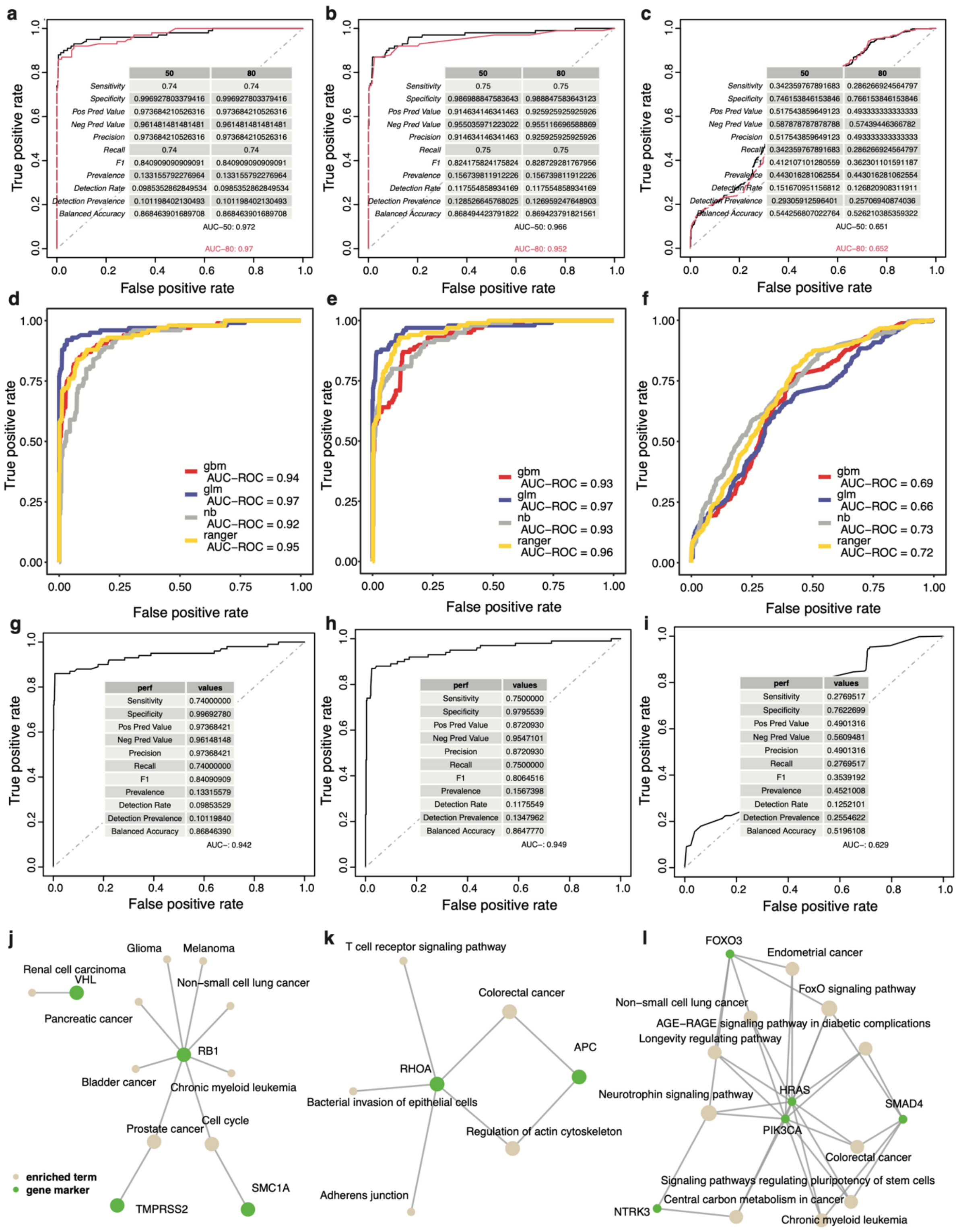


**Supplementary Figure S6. Discovery of genomic events linked with the EMT macro-states.** (a-c) ROC curves and statistics for the lasso model predictions based on genomic markers distinguishing between MES and EPI (a), MES and hEMT (b), or hEMT and EPI (c) states, respectively. (d-f) ROC curves for predictions based on genomic markers obtained with the lasso procedure and tested using different machine learning approaches, for the MES versus EPI (d), MES and hEMT (e), or hEMT and EPI (f) models, respectively. (g-h) Same as (a-c) but for random forest based models. (j-l) Pathway enrichment for gene events significantly distinguishing the the MES from the EPI group (j), the MES from the hEMT group (k) and the hEMT from the EPI group (l).

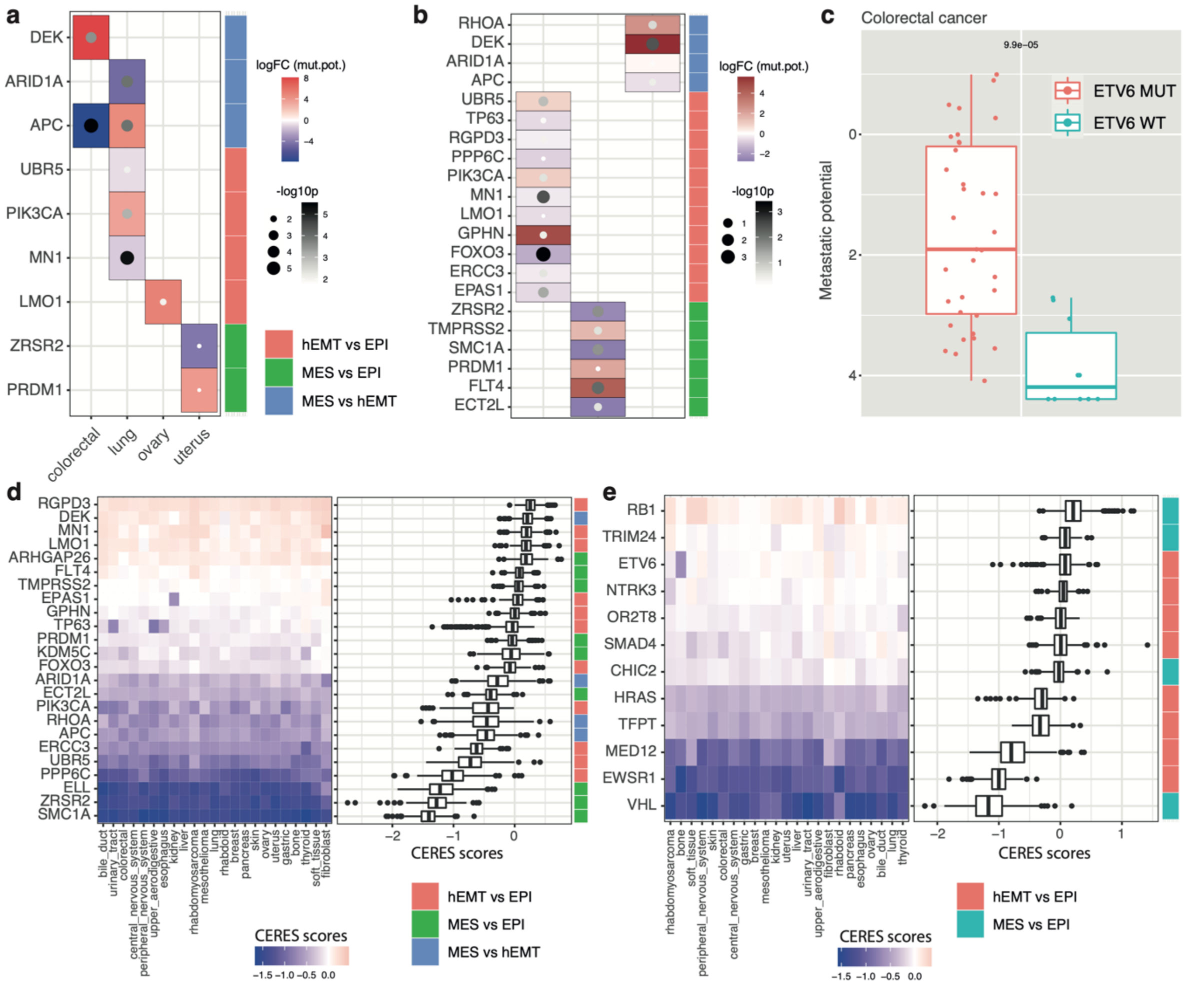


**Supplementary Figure S7. Validation of the pan-cancer genomic associations with EMT in external datasets.** (a) Fold changes in metastatic potential across cell lines from CCLE originating from different tissues and harbouring copy number alterations in marker genes of EMT, compared to that of cell lines without the respective alteration. The size and the colours of the dots highlight the significance of the association ($p<0.05$). (b) Similar to (a), but with the analysis performed pan-cancer rather than at tissue level, ($p<0.05$). (c) An increase in metastatic potential is observed in colorectal cancer cell lines harbouring an

ETV6 mutation. (d) CERES essentiality scores from DepMap in individual cell lineages for genes harbouring copy number alterations linked with EMT. Negative values indicate increased essentiality. The boxplots on the right indicate the CERES score distribution across all lineages. (e) Similar to (d) but considering genes harbouring point mutations.

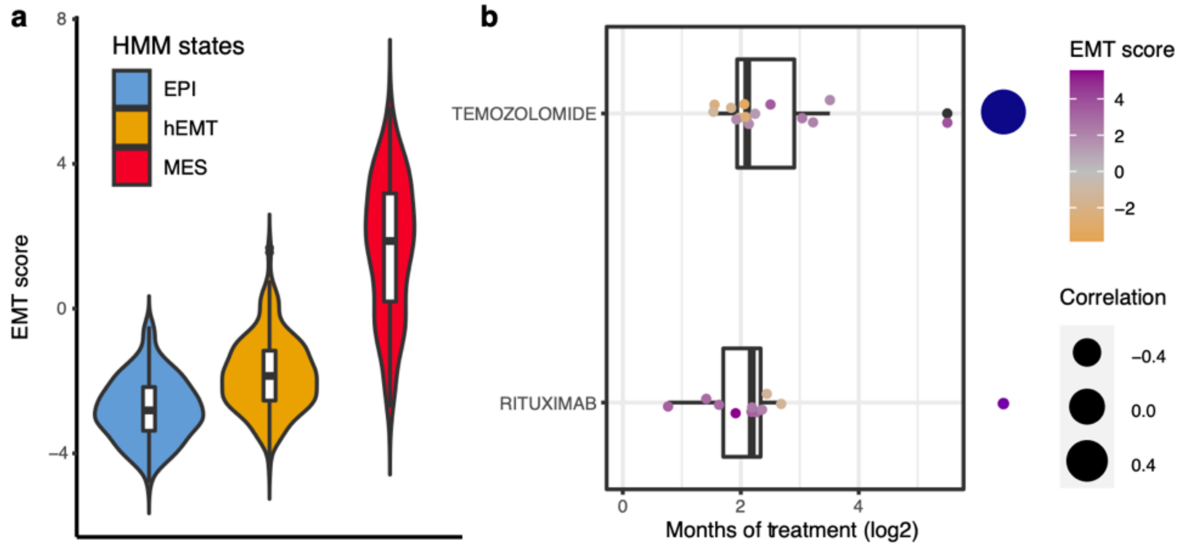


**Supplementary Figure S8. Therapeutic relevance of the EMT states in the POG570 cohort.** (a) EMT scores are compared across distinct macro-states following EMT reconstruction of the post-treated samples from POG570. The differences observed confirm the classification. (b) Duration of drug treatment is correlated with an EMT score increase (temozolomide) and decrease (rituximab) for selected drugs. Only significant associations are shown.

### SUPPLEMENTARY TABLE CAPTIONS

**Supplementary Table S1. Pseudotime reconstruction of EMT trajectories in TCGA samples.** The results of the pan-cancer reconstruction of epithelial-to-mesenchymal trajectories is reported, along with the HMM state of the sample, and the corresponding EMT macro-state and score.

**Supplementary Table S2.** Distribution of the TCGA samples in each EMT state across clinical cancer stages.

**Supplementary Table S3.** Stratification of samples according to hypoxia score and CD44 expression.

**Supplementary Table S4**. **Pan-cancer genomic events linked with EMT states.** The results of the lasso models built to distinguish EMT states based on genomic markers are shown. All genes included in at least 50% of the models are listed. The first column indicates the comparison in which a genomic marker (second column) has been identified. Additional details on the genes are provided. The mean coefficients from the lasso model (Mean contribution). The final column annotates the genes comprised within an altered chromosomal arm.

**Supplementary Table S5**. **Putative EMT biomarkers and literature evidence for links with EMT.** Genes included in at least 80% of the lasso models are listed, along with the model(s) in which they were included and PubMed IDs (PMID) of publications where they are linked with EMT, cell migration or cancer progression. There was weak or no literature evidence for EMT associations for the genes highlighted in red.

**Supplementary Table S6. Cox regression model results on clinical end points.** (a) Results of the multivariate Cox regression model for the overall survival. The mean and standard deviation are reported for each covariate along with the hazard ratios (HR) from the univariable and multivariable models. The p-value for the model is highly significant ($p<1e-114$). (b) Results of the multivariate Cox regression model for the progression free interval

(PFI). The mean and standard deviation are reported for each covariate along with the hazard ratios (HR) from the univariable and multivariable models. The p-value for the model is highly significant (p<1e-79).

**Supplementary Table S7. Mutation events impacting overall survival in TCGA.** The first column indicates the EMT comparison model the marker was derived from, the second column reports the name of the gene. Hazard ratios (log10) are reported along with confidence intervals, p-values and outcome (Out: 'positive' indicates better prognosis for patients harbouring the mutation in the respective gene, 'negative' indicates worse prognosis).
